# Horizontal orientation facilitates pollinator attraction and rain avoidance in radially symmetrical flowers

**DOI:** 10.1101/2021.09.08.459386

**Authors:** Taichi Nakata, Ishii Rin, Yuki A Yaida, Atushi Ushimaru

## Abstract

**Premise:** Floral angle, such as upward, horizontal, and downward orientation are known to evolve under both biotic and abiotic agents to enhance pollination success in zoophilious plants. Adaptive significance of horizontal orientation in radially symmetrical (actinomorphic) flowers under biotic and abiotic selection pressures were largely unknown, although those in bilaterally symmetrical flowers have been well studied.

**Methods:** Using experimentally angle changed flowers, we examined the effects of flower angle on pollinator behaviors, pollination success and rain avoidance in a population of insect-pollinated *Platycodon grandiflorus*. We further investigated the frequency and amount of precipitation in the flowering season and pollen damage by water in this species. *Main results*: Horizontally oriented flowers received more visitations and pollen grains on the stigma in male and/or female phases than downward and/or upward oriented flowers and avoided pollen damage by rainfall compared to upward oriented flowers. The pollen germination experiment showed that approximately 30% of pollen grains burst in distilled water, thus pollen damage by rainfall was potentially serious in *P. garndiflorus*.

**Conclusion:** In this study, our field experiments revealed that upward flowers cannot avoid damage from rainfall during the flowering period whereas both upward and downward flowers suffered from pollinator limitation in female success. Thus, horizontal flower orientation is suggested to be adaptive in this insect-pollinated actinomorphic species which blooms in the rainy season.

## Introduction

Present floral diversity in angiosperms are considered to have evolved under selections mediated by both biotic and abiotic agents (Darwin, 1862; Grant and Grant, 1965; Stebbins, 1970; Fenster et al., 2004; Wilmer, 2011). In animal-pollinated species, most floral traits such as size, shape, color, scent, flowering timing have adapted to enhance pollen transfer by their respective pollinators, while some of the traits simultaneously function as protection against harmful abiotic factors such as rain and very low/high temperature (Kudo, 1995; Huang et al., 2002; Patino et al., 2002). Flower angle (e.g. vertical direction of flower orientation including upward, horizontal, oblique and downward orientation) regulated by flower stalk angle is a trait which evolves under both biotic and abiotic agents to enhance pollination success in zoophilous plants (Hocking and Sharplin, 1965; Kevan, 1975; Kudo, 1995; Tadey and Aizen, 2001; Huang et al., 2002; Patino et al., 2002; Galen and Stanton, 2003; Ushimaru et al., 2009; Haverkamp et al., 2019).

Flower angle is known to influence attraction to and behavioral control of specialized pollinators (Fenster et al., 2009; Ushimaru and Hyodo, 2005). Bilaterally symmetrical (zygomorphic) flowers usually exhibit horizontal orientation (Neal et al., 1998), which can attract more pollinators and enhance their legitimate behaviors for achieving higher pollen transfer success compared to upward or downward orientation (Ushimaru and Hyodo, 2005; Ushimaru et al., 2009; Wang et al., 2014a). In contrast, the optimal flower angle may vary among radially symmetrical (actinomorphic) flowers likely depending on pollinator composition. For example, upward and heliotropic orientation increases floral temperature to attract many fly pollinators in alpine, arctic and early-spring blooming plants with dish-shaped actinomorphic flowers (Hocking and Sharplin, 1965; Kevan, 1975; Kudo, 1995). Downward orientation of tubulous actinomorphic flowers likely limits pollinators to some specialized groups such as humming birds and large bees, improving pollination efficiency whereas upward-orientation facilitate nocturnal hawkmoth pollination in actinomorphic species (Fulton and Hodges, 1999; Aizen, 2003; Campbell et al., 2016).

Meanwhile, pollinators are unlikely the primary selective agent driving the evolution of floral angle but abiotic factors put more severe pressures in some zoophilous plants (Haung et al., 2002; Tadey and Aizen, 2001; Wang et al., 2010; Lin and Forrest, 2019). Downward flower orientation is thought to have evolved to avoid pollen damage and nectar dilution by rainfall and exposure to solar radiation in actinomorphic flowers (Huang et al., 2002; Tadey and Aizen, 2001; Wang et al., 2010; Lin and Forrest, 2019). In some generalist actinomorphic flowers which are pollinated by a wider range of pollinator groups (two or more functional groups), flower angle usually less influences pollinator composition and visit frequency, so that the effects of rainfall might be a more important driving force selecting downward orientation (Huang et al., 2002; Wang et al., 2010; Lin and Forrest, 2019).

The adaptive significances of horizontal orientation in actinomorphic flowers under biotic and abiotic selection pressures have not been examined to date, although those in zygomorphic flowers are relatively well studied. Generally, specialized zygomorphic flowers exhibit higher pollination success when flowers face horizontally (Neal et al., 1998; Ushimaru et al., 2009; Wang et al., 2014a; Armbruster and Muchhala, 2020). So far only a single study examined the role of horizontal orientation in a generalist zygomorphic species, in which horizontal oriented flowers achieved higher pollen transfer success than upward and downward orientated flowers (Yu et al., 2020). In the same species, horizontal flower orientation could reduce pollen damage by rainwater compared to upward orientated flowers, but the anthers were always wet during rainfall irrespective of flower angle (Yu et al., 2020). These together indicate that horizontal orientation have mainly evolved to attract and control pollinators in zygomorphic flowers. Meanwhile, to the best of our knowledge, functional roles of horizontal orientation in actinomorphic flowers with generalist systems were still unknown. We may expect that pollinator-mediated selection on flower angle would be less strong in generalist actinomorphic flowers.

Here, we report a case study on functional roles of horizontal orientation in attraction and landing control of pollinators, pollen transfer and rainfall avoidance in actinomorphic *Platycodon grandiflorus* whose flowers are visited by diverse insect groups in the rainy season. We conducted a flower angle-change experiment to examine the effects of flower angle on pollinator behaviors and pollination success. We also examined pollen viability in water and sucrose solutions and precipitation during the flowering season in the study site to investigate possibility of pollen damage by rainfall. Flowers of *P. grandiflorus* are protandrous and male and female phases do not overlap temporally within each flower. We therefore examined and discussed whether there were difference in the effects of flower angle between male and female phases in this study.

## MATERIALS AND METHODS

### Study species and site

The Japanese bellflower, *Platycodon grandiflorus* (Jacq.) A.DC. (Campanulaceae) is a perennial herb species, which is currently listed as Vulnerable species in the Natural Red list (Ministry of the Environment of Japan, 2020). This species usually grows on natural and semi-natural grasslands in Japan, Korean Peninsula, China and east Russia. In Japan, the species blooms during the rainy season from mid-June to mid-September. Blue cup-shaped insect-pollinated flowers, which are usually open for 4–6 d, are self-compatible and protandrous and require pollination for fruiting and seeding (Wei et al., 2006): male and female phases did not overlap and lasted average 1.3 d (1–3 d, n = 170) and 2.9 d (2–5 d, n =132), respectively (Fig. S1b, c). Both in male and female phases, flowers oriented nearly horizontally: flower angles deviated 2.4° in average (−43–32°, n = 32) from the horizontal direction (Fig. S2). In a male phase flower, the pistil produces pollen-bearing hairs, on which secondary pollen presentation occurs soon after flower opening whereas female phase starts with curling the stigmatic lobes like in other Campanulaceae species (Vranken et al., 2014) More than 80% of pollen grains have viability just after flower opening and pollen vigor quickly decreased to less than 30% three days later in cultivated plants (Wei et al., 2006). An individual of this species has thick roots and usually does not spread clonally.

We examined a population of *P. grandiflorus* in the Sugadaira plateau, Nagano Prefecture, Japan (36°32’12.9 N, 138°20’53.1 E) in August 2018–2019, 2021. The species is distributed only on ski slope grasslands which are managed by annual mowing in early September (Yaida et al., 2019). In the study site, we observed several pollinator groups visiting *P. grandiflorus* flowers to forage pollen and nectar: large and medium sized bees (LM bees, mostly *Megachile* bees and more infrequently *Bombus* and *Apis* bees), small-sized bees (S bees, such as *Andrena* and Halictidae species), shyrphid flies (*Episyrphus balteatus, Syrphus torvus* and other syrphid flies), infrequently scolid wasps and butterflies and skippers (e.g., *Ochlodes ochraceus*) and beetles (Fig. S3).

### Flower diameter change during opening

We measured flower diameters of en face surface at male (ca. 0.5-1d after opening) and female phases (over 3 d after opening) by a digital caliper (in mm) to examine the effects of temporal change in petal lobe opening on pollinator attraction and rainfall avoidance.

### Effects of floral angle on pollinator behaviors

We experimentally prepared three types of flowers that differed in terms of their floral angle in 2018:

1. Con: control flowers whose angle were not changed,
2. Up: flowers whose faces were turned upright and
3. Down: flowers whose faces were turned downward.

Angles of all experimental flowers were fixed by short wires during the anthesis (Fig. 1). The same treatment was done for all opening flowers within the same individual: totally 413 (143 Con, 139 Up and 131 Down) flowers on 84 (31 Con, 28 Up and 25 Down) individuals: four flowers on the same plant were examined at the maximum. The number of opening flowers for each individual varied 1 to 3 and the average was 1.4 in the site. All the experimental individuals and their flowers were marked and identified.

**FIGURE 1.**
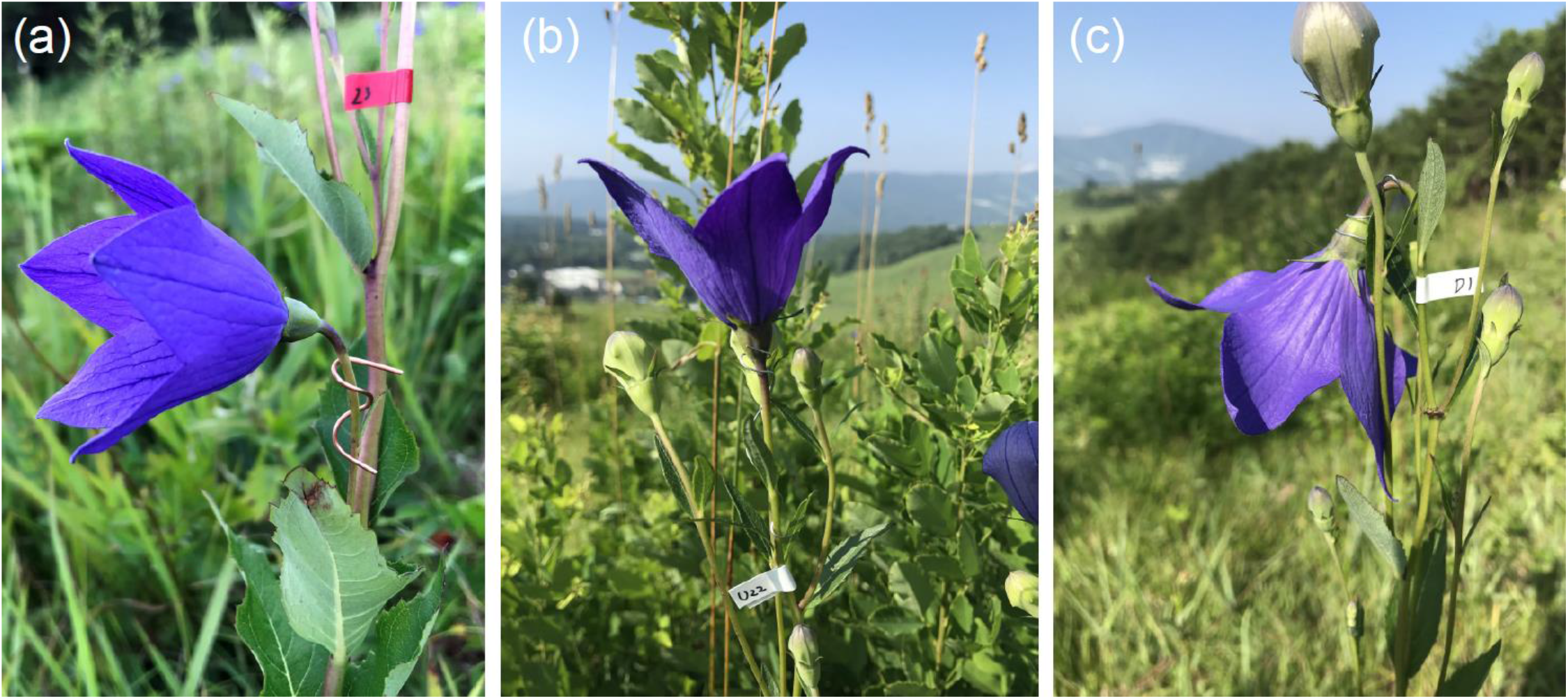
Photos of experimental flowers of *Platycodon grandiflorus*: control (Con, a), upward-facing (Up, b), and downward-facing (Down, c) flowers. Main floral axes of Con and manipulated (Up and Down) flowers were prepared nearly horizontal and vertical, respectively.

To test the effects of floral angle on pollinator behaviors, we observed pollinator visits to experimental flowers. For each trial, we arbitrarily selected neighboring 3–10 experimental individuals including more than one flower types and observed pollinators to them for 20-min observation. We conducted totally 41 observation trials (820 min observation in total) during the daytime from 08:00 to 15:00 h on sunny days when pollinators were active. Before each trial, we recorded sexual phase (male or female) of each experimental flower. Most flowers were repeatedly observed in different timings during the same sexual phase in the same day and/or on different dates and at different sexual phases: the average number of observation times per flower was 3.36 (1–8) and 16 flowers were observed only once. Twenty-five and 56 flowers were observed in either male or female phase, respectively whereas 42 flowers were observed in both male and female phases. In all trials, we observed cumulatively 95 male Con, 87 male Up, 97 male Down, 48 female Con, 52 female Up and 34 female Down flowers.

During our observation, we recorded two types of pollinator behavior: approach and landing behaviors. Approach occurred when pollinators found a flower and approached it from the front. Landing was defined as a pollinator landing on any part of a flower after approaching it. We further divided landings into the following two types (Ushimaru and Hyodo, 2005): legitimate landing, a pollinator touched any parts of the pistil and stamens during its landing; petal landing, a pollinator landed on any part of petals and foraged or collected pollen without touching secondary presented pollen on the pistil or the stigma lobes. We counted the number of each behavior on each experimental flower for each observation trial.

### Effects of floral angle on pollen transfer success

We estimated pollen removal by counting the number of pollen grains remaining on the pistil hairs of 60 experimental flowers and that on each of newly opened, unvisited ten flowers in 2019 (c.f.; Harder 1990; Ushimaru et al. 2006; Ushimaru et al. 2014; Katsuhara et al. 2017; Ushimaru et al. 2021). We collected a single pistil form each of naturally-pollinated 20 Con, 20 Up and 20 Down flowers (60 individuals) whose male phases were almost finished by carefully checking the pistil conditions, such as stigmatic lobes status, and stored separately. The pistil with secondary pollen presentation were collected from each of ten unvisited flowers after their opening and was stored in 1.0 mL 99.9 % ethanol as a single sample. We vortexed the sample well and estimated the number of pollen grains per sample by counting the pollen numbers in three 10.0 μL drops per sample under a microscope. To quantify pollen removal from each experimental flower we calculated pollen removal [(the mean estimated number of pollen grains in the newly opened flowers) - (the estimated number of pollen grains remaining in each experimental flower)] as an integer (Ushimaru et al., 2014; Ushimaru et al., 2021).

We also examined the effects of floral angle on pollen receipt by collecting the pistil of a single flower form each of 20 Con, 20 Up and 20 Down individuals ca. 3 d after the start of female phases in early August, 2019. These flowers were exposed to natural pollinator visits from bud break for ca. 5 d. We dissected the stigma from a remaining part of each pistil and stored it separately. We then counted the number of pollen grains deposited on the stigma for each sampled flower under a microscope (× 40) in the laboratory.

### Experimental flowers in the rain and precipitation during the flowering season

In 2019, though it was not quantitatively, we observed several male-phase and female-phase experimental and intact flowers after rainfall to examine whether mating related parts (anthers, pistil hairs and stigma) were wetted, soaked or not.

Additionally, to examine the effect of flower angle on rain susceptibility, 61 experimental flowers (10 male Con, 10 male Up, 10 male Down, 11 female Con, 10 female Up, 10 female Down) were prepared and observed whether flower base, mating related organs and pollen of these flowers were wet or soaked 24-h after rain in 16 August, 2021.

We obtained meteorological data during the last decade (2010–2019) from the nearest AMEDAS (governmental weather observation system in Japan; 36°31’9 N, 138°19’5 E) station to examine precipitation (mm per day) during the flowering season (15 July–15 September) of *P. grandiflorus* in the study site.

### Pollen germination in water and sucrose solution

To test how rainfall influence viability of pollen grains, we examined pollen germination and burst in water or sucrose solutions according to the existing method (Dafni, 1992; Huang et al., 2002). We collected pollen grains from 10 flowers just after opening and preserved at ca. -20 °C in a freezer in our laboratory. We then placed pollen grains from dehisced anthers of a single flower on slide glasses with 0, 5, 10, 15, and 20 % (g/g × 100) sucrose solutions: the 0% solution is just distilled water. For each glass, we examined the fates of 45–235 (average 145.7) pollen grains and counted the number of germinated and burst pollen grains after 24 h under a light microscope. This germination/burst test was repeated ten times for each sucrose concentration.

### Analyses

We first compared en face flower diameter between sexual phases using a generalized linear mixed model (GLMM, gaussian errors and identity link function) in which the diameter of each flower, sexual phase (male/female) and individual identities were the response and explanatory variables and the random terms, respectively. We then examined the effect of angle changes on pollinator behaviors using GLMMs with Poisson errors and logarithmic link function. The number of approaches, legitimate landings or petal landings per 20 min. per flower was the response variable. In all the models, we included treatment (Con/Up/Down), sexual phase and their interaction as the explanatory variables, display size (the number of flowers per individual) and observation time (start time, minutes from 08:00) as covariates and the individual identity as a random term. We also conducted the same GLMM analyses for the two behavior of three major pollinator groups, separately. Secondly, we compared pollen removal and receipt between Con and other (Up and Down) flowers using generalized linear models (GLMs with negative binomial errors and logarithmic link). The model incorporated treatment as the explanatory variable and pollen removal or pollen receipt was the response variable. Thirdly, we examined difference in the ratio of flowers whose flower base or mating related organs among experimental flowers using a Fisher’s exact test. We finally compared the ratios of germinated and burst pollen grains (burst or germination) on using GLMMs (negative binomial error and logarithmic link), in which each pollen number, the sucrose concentration category (0/5/10/15/20), and total number of examined pollen grains were the response and explanatory variables, an offset term, and individual identity as a random term respectively. All statistical analyses were done using the software R (R Development Core Team 2015).

## RESULTS

### Flower diameter change

We found a significant increase in en face flower diameter from male (mean ± SE, range; 5.0 ± 0.24 cm, 2.0–5.1 cm) to female (5.8 ± 0.49 cm, 2.0–7.3 cm) phases (Table S1), indicating that petal lobes opened more and often bent backward in the female phase of each flower. Note that the whole style was under the umbrella of upper petal lobe during the male phase in the horizontally orientated flowers (Fig. 1a).

### Effects of floral angle on pollinator behaviors

Most dominant pollinators to experimental flowers were LM bees (227 approaches, 56.7 % of total approaches) and S bees (100 approaches, 24.9 %) and syrphid flies (74 approaches, 18.5 %) followed. It should be noted that the ratios of the three major pollinator groups did not differ significantly among flower types and between sexual phases for legitimate landings (Fig. S4b).

Approach frequency per flower by all pollinators did not differ significantly between Con and other flower types whereas approaches to male-phase flowers were significantly higher than those to female-phase flowers in all the flower types (Fig. 2a, Table S1). Approaches per flower by all pollinators significantly decreased with increasing display size (Table S1). Significant decreases in approach at female phase and with larger display size were found in LM and S bees, respectively. Approaches significantly increased with observation time in all pollinators and bee groups (Table S1).

**FIGURE 2.**
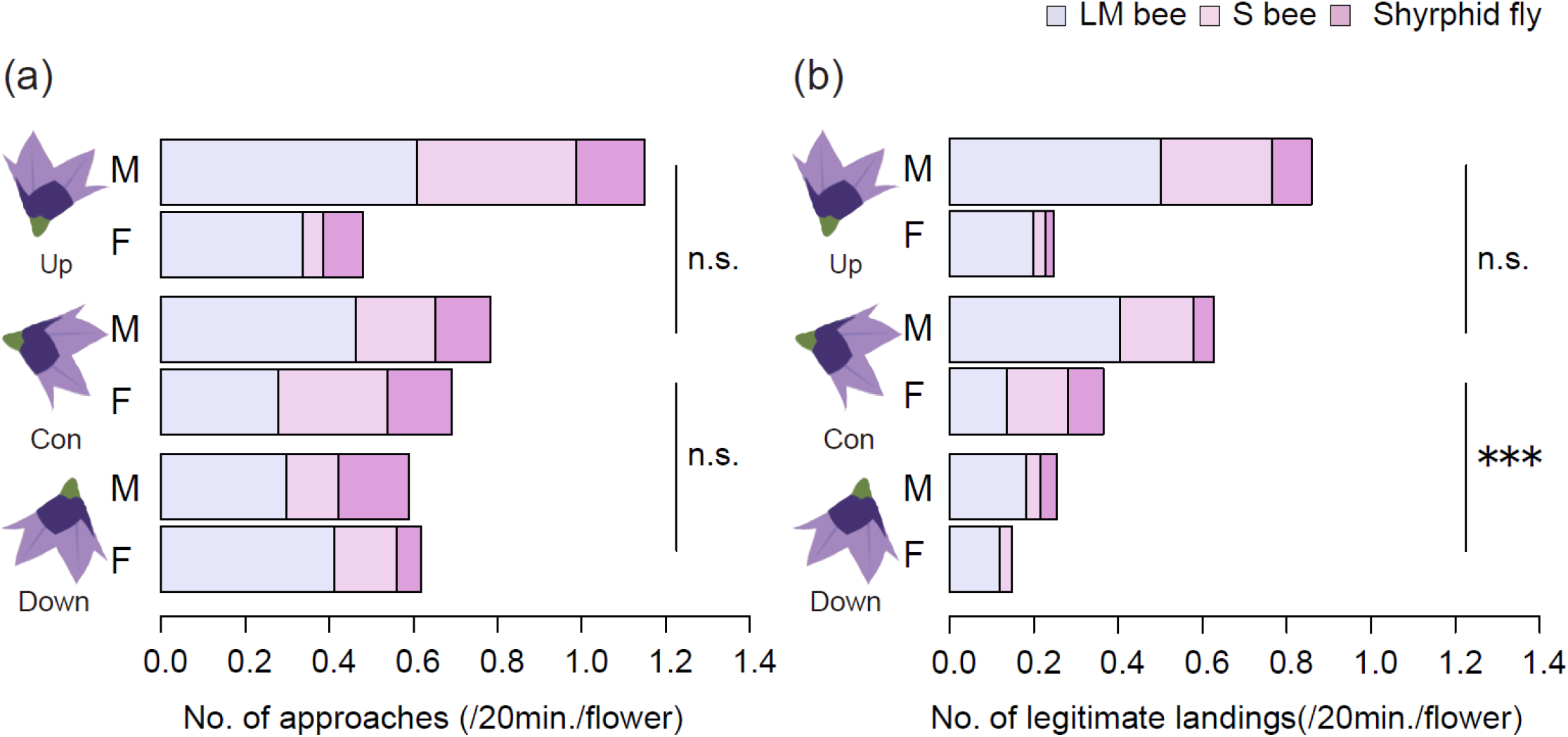
Mean numbers of approaches (a) and legitimate landings (b) by pollinators per 20 min per flower for the experimental (Con, Up, and Down) flowers of *Platycodon grandiflorus*. M and F show each sexual phase (M: male phase; F: female phase). ***, *P* < 0.001; n.s., *P* > 0.1, by GLMM.

Legitimate landings by all pollinators and bees were significantly fewer on Down flowers than on controls whereas the variable did not differ between Con and Up flowers (Fig. 2b, Table S1). The response variable for all pollinators and LM bees was significantly lower in female phase flowers than in male-phase flowers (Fig. 2b, Table S1). Legitimate landings increased with observation time in all pollinators and bees whereas the variable decreased with increasing display size only in S bees (Table S1).

Petal landings by all pollinators and each pollinator group was not influenced by any explanatory variables at all (Table S1). The interaction between flower angle and sexual phase had no significant effects on any response variables in any pollinator groups except for approaches by S bees, such that the variable significantly increased in female-phase Up flowers than male-phase controls (Table S1).

### Effects of floral angle on pollen transfer success

More than 95 percent of pollen grain were removed from pistil hairs in all the flower types and significant higher pollen removal was found in Up and Down flowers than in controls (Fig. 3a, Table S1). By contrast, the stigmas of Up and Down flowers received significantly fewer pollen grains (ca. 45 and 30 grains in average, respectively) than those of Con flowers which received more than 65 pollen grains in average (Fig. 3b, Table S1).

**FIGURE 3.**
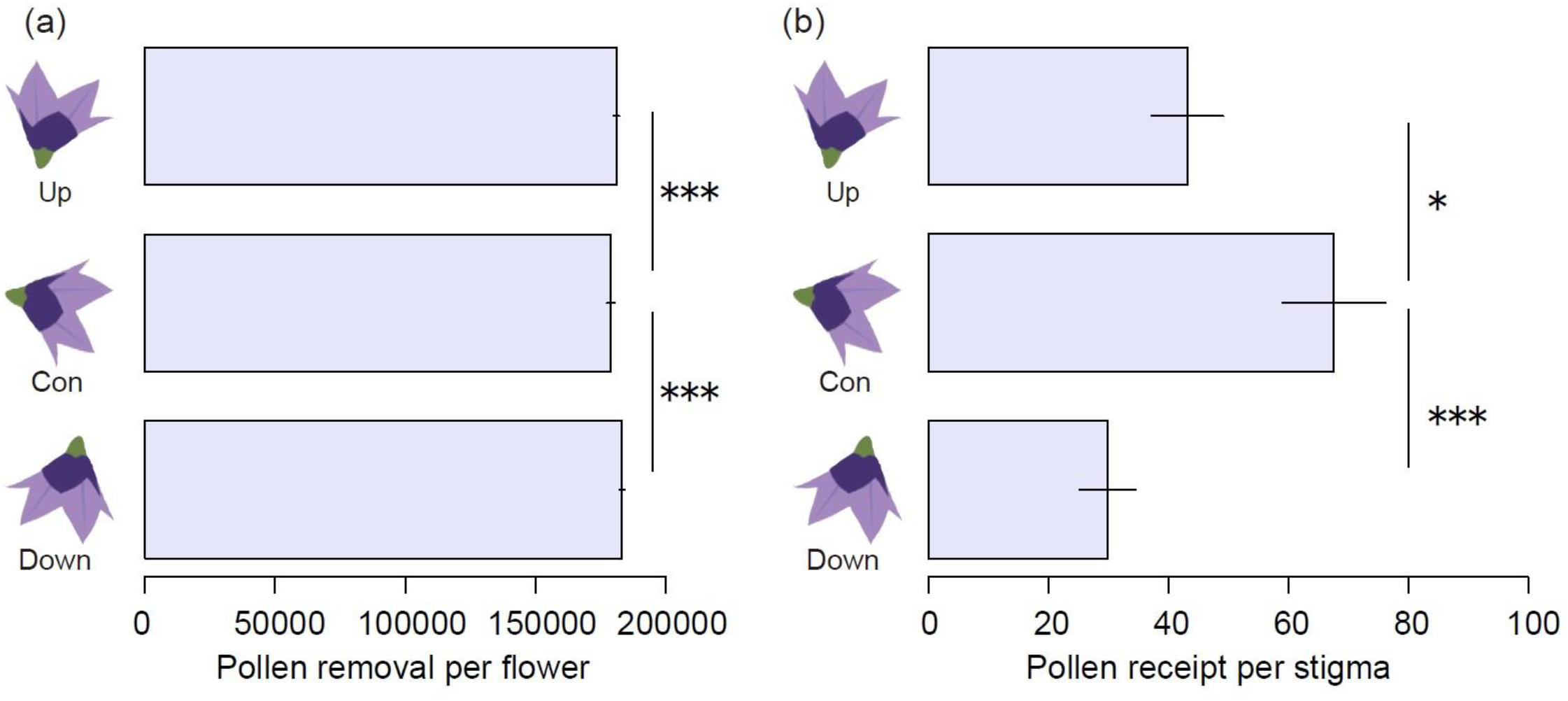
Mean number of remaining pollen grains on the pistil hairs (a) and deposited on the stigma (b) for the experimental (Con, Up, and Down) flowers. Bars show standard errors. ***, *P* < 0.001; *, *P* < 0.05 by GLM.

### Flowers in rainy conditions and precipitation during the flowering season

When over 3.5 mm precipitation per day was observed, Up flowers accumulated rainfall and the anthers and pistil hairs therein were soaked in the male phase whereas some parts of the stigma lobes of Up flowers were wet or soaked in the female phase in 2019 (Fig. S5a, b). In Down flowers, mating related parts were never wet or soaked, although the back of petals were wet. The back of upper petals and the front of lower petal lobes were wet in Con and intact flowers, but the mating-related organs were rarely wet: stigma surfaces were very infrequently wet in female-phase flowers with very opened petal lobes.

The flower base was soaked or wet after 6.0 mm rain in many Up flowers in 16 August, 2021, although not much water remained in flower cups owing to stem oscillation by the strong wind. We observed a single case in which the pistil of Up flower was broken by the rain (Fig. S5c). The number of flowers whose pollen, flower base and/or mating related organs were soaked or wet was significantly greater in Up flowers than in other flower types (Fig.4a). In Con and Down flowers, lobes and backs of the petals were often wet but mating related organs were never wet in the experiment (Fig. 4b).

**FIGURE 4.**
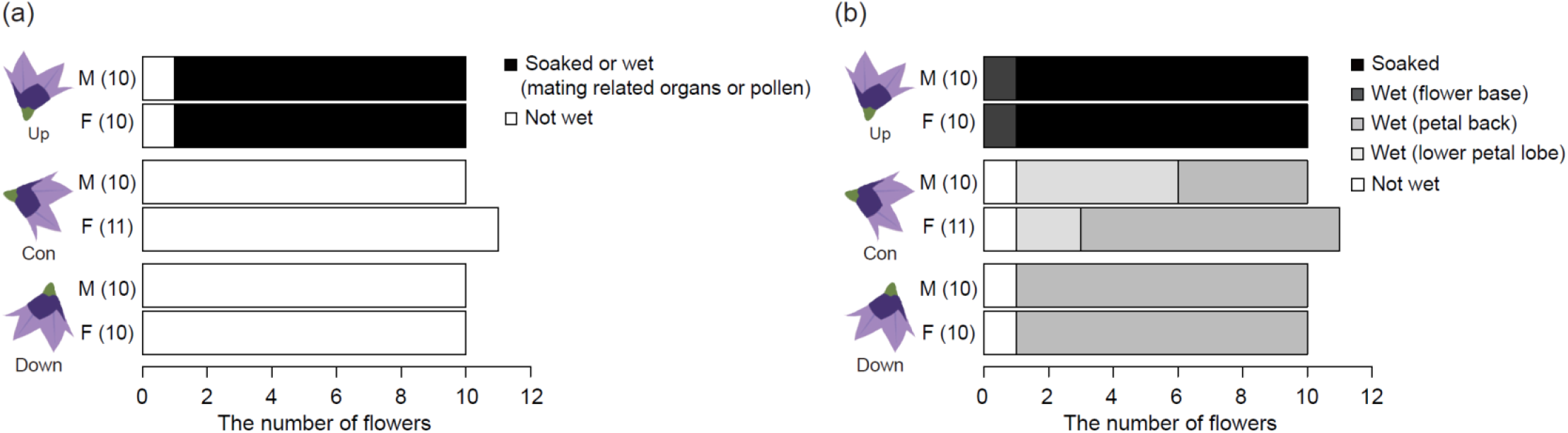
The effect of flower angle on rain susceptibility in experimental *Platycodon grandifloras* flowers. The ratios of flowers whose flower base (a) and mating related organs or pollen (b) were soaked or wet after ca. 6.0 mm rainfall differed significantly among flower types (a, b; Fisher’s exact test, *p* < 0.001).

In the last decade, ≥ 3.5mm precipitations, which was enough to soak most part of the pistil in Up flowers, were found for 10–21 (mean, 16.6) d during the flowering period (Fig. S6). Moreover, heavy rain over 10 mm per hour were often observed in the study area. More than 4.2 sequential days (the average longevity of a flower) without rainfall were observed less than half of the flowering season in 2010–2019, indicating that the majority of flowers experienced rainfall to some extent during their opening in the study area (Fig. S6). The number of periods when more than 5 sequential days without rain were only 3 times and once in 2018 and 2019, respectively in the study areas during the flowering period (Fig. S6).

### Pollen germination in water and sucrose solution

On average, 56.3 % of pollen grains germinated in the 5–20 % sucrose solution whereas only 0.04 % of grains germinated in distilled water (Fig. 5). The differences between distilled water and sucrose solutions were significant (Table S1). Approximately 30 % of pollen grains burst in distilled water. By contrast, only less than 0.007 % of pollen grains burst in 5–20 % sucrose solutions and the percentages were significantly lower than that in distilled water: especially no pollen burst was observed in 20 % sucrose solutions.

**FIGURE 5.**
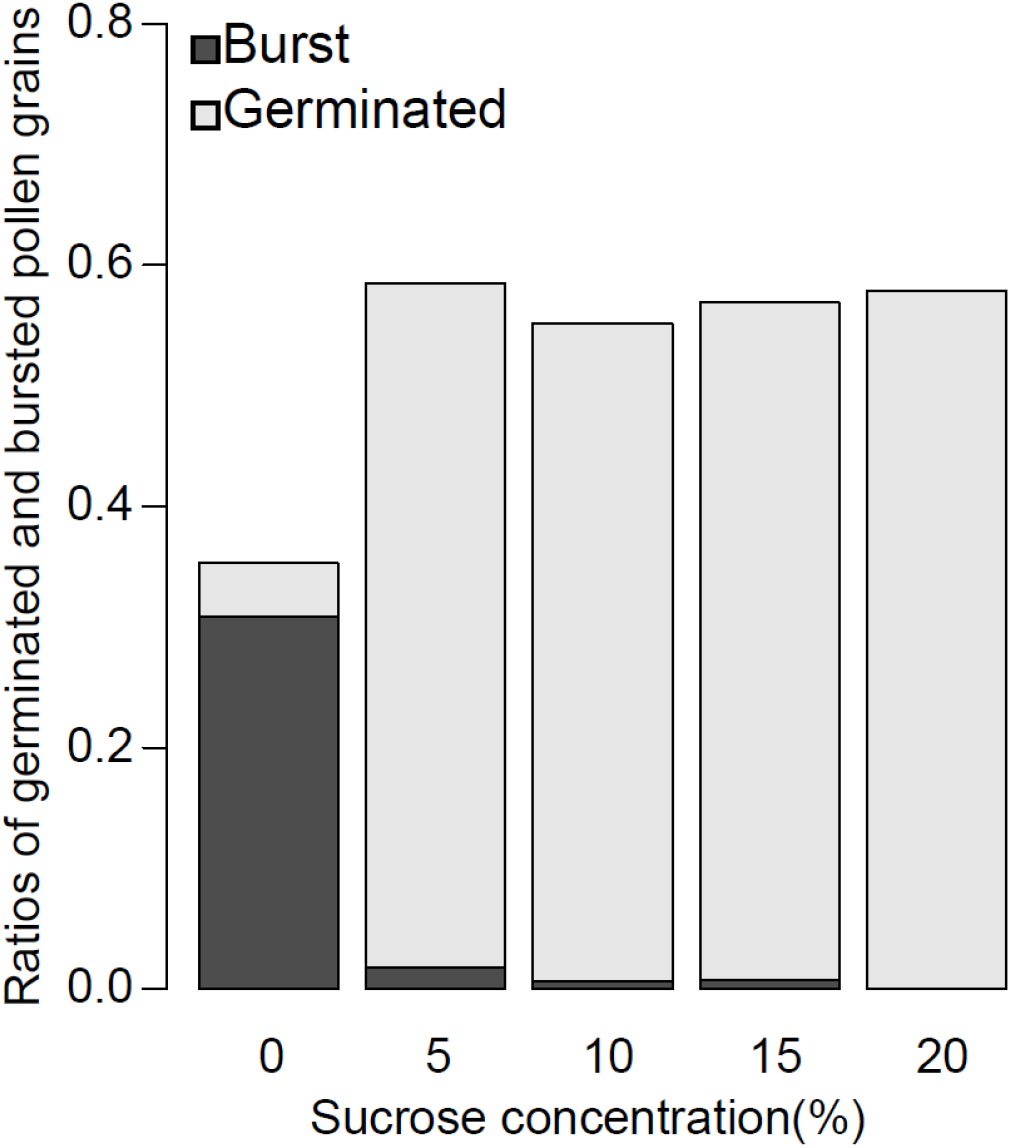
Ratios of germinated and burst pollen grains of *Platycodon grandifloras* in sucrose solution with various concentrations.

## DISCUSSION

In this study, we examined functional roles of horizontal flower orientation with respect to pollinator behavior control and rain protection in actinomorphic *P. grandiflorus*. Our filed experiments showed that upward oriented flowers likely suffered more from pollen and pistil damages by rainfall and pollen limitation than controls whereas downward oriented flowers received less visitations and pollen grains on the stigmas compared to control flowers. Thus, horizontal flower orientation might have evolved under both pollinator- and rain-mediated selections in actinomorphic *P. grandiflorus* flowers. We discussed our findings more in detail below.

### Effects of floral angle on pollinator behaviors and pollen transfer

Although all flower types experienced a similar number of approaches by all pollinator groups, downward flower orientation reduced legitimate landings of bees when comparing to horizontal and upward orientation in both male and female phases unlike our expectation. Thus, floral orientation influenced pollinator landing behavior rather than pollinator attraction. The results are concordant with the findings in zygomorphic *Commelina communis* (Ushimaru and Hyodo 2005), but not with patterns in another zygomorphic species, *Corydaris sheareri* whose upward-oriented flowers also had pollinator limitation (Wang et al., 2014a) as well as in actinomorphic flowers of *Pulsatilla cernua* (Haung et al., 2002), *Geranium refractum* (Wang et al. 2014b) and *Mertensia* species (Lin et al., 2019) whose downward-oriented flowers did not suffer from pollinator limitation. The discrepancies between the present and previous studies (Wang et al., 2014a, b; Lin et al., 2019) might be owing to differences in pollination system, i.e., a set of floral characteristics and pollinators. Bumble bees usually less discriminate downward flowers with open or cup shapes but leafcutter and small-sized bees, syrphid flies and lepidopterans do when approaching and/or landing (Haung et al., 2002; Ushimaru and Hyodo 2005; Wang et al., 2014a, b; Lin et al., 2019; Yu et al., 2020). Moreover, long-tubed flowers were often not preferred by diurnal pollinators when facing upright experimentally (Wang et al., 2014a; Yu et al., 2020) whereas upright open or cup-shaped flowers were visited by various groups of pollinators who usually used petals as landing platforms (Huang et al., 2002; Ushimaru and Hyodo 2005; Wang et al. 2014b; Lin et al., 2019). Leafcutter bees, small-sized bees and syrphid flies (the dominant pollinators in *P. grandiflorus*) were sometimes observed to take longer time to find footholds on Down flowers than on other flower types, although we did not quantify the time. The longer handling time might cause discrimination of Down flowers by these pollinators.

Pollen removal per flower was significantly enhanced by upward and downward flower orientations compared to horizontal one. Though the difference was not significant, upright flowers experienced more legitimate landings than controls in the male phase, being consistent with the pollen removal result. Meanwhile, pollinator-limited downward flowers had higher pollen removal than controls did as well. We cannot explain this with our data set, but accidental pollen loss owing to shaking flowers by pollinators and wind could be responsible. Anyway, over 95 % of pollen grains were removed from all flower types, so the difference in pollen removal was small (Fig. 3).

By contrast, flower angle change treatments largely reduced pollen receipt on the stigma in *P. grandiflorus* like in the previous studies on horizontally-oriented zygomorphic flowers (Ushimaru et al., 2009; Wang et al., 2014a). Pollinator limitation might cause lower pollen receipt in downward flowers. Upward flowers received lower numbers of visits than control flowers in the female phase though not significant (Fig. 2a), likely causing lower pollen receipt. However, further clarification why upward flowers had lower female pollination success would be required in future.

Sexual phase, display size and time in the day influenced pollinator visitation to *P. grandiflorus* flowers. Female-phase flowers received significantly less approaches and landings by larger-sized bees than male phase flowers, suggesting that these bees foraged mainly pollen on flowers. Larger display sizes tended to decrease pollinator attraction per flower, indicating that relatively large corolla of each flower enough attract pollinators and that simultaneous opening of multiple flowers within an individual might be rather negative for reproductive success of individual flowers. This may explain relatively small average display size (1.3 flowers per plant) of this species. Bees increased their visitations to flowers toward noon and afternoon but no such a pattern was observed for syrphid flies. Daily active patterns are known to vary between bees and flies (Herrera, 1990; Rader et al., 2013; Ushimaru et al., 2021). However, to elucidate factors explaining the difference, more information on thermoregulatory ability, energy requirements and their interactions with habitat environments for each group (or species) were needed (Herrera, 1990; Rader et al., 2013)

### Role of horizontal flower orientation in rain avoidance

It is well known that pendant and downward flowers have a function to protect pollen grains from rainfall and sun radiation (Tadey and Aizen, 2001; Haung et al., 2002; Haverkamp et al., 2019). In *P. grandiflorus,* many pollen grains burst and lost the germination ability in water, indicating a significant negative effect of rainfall on the pollen viability like in the other species (Tadey and Aizen, 2001; Huang et al., 2002; Mao and Haung, 2009). The anthers, pistil hairs and stigma of control flowers were observed not to be wet and soaked as in experimental downward flowers (Fig. 4, S5d, e, f, g). Interestingly, in the male phase, petal lobes did not fully open compared to those in the female phase, suggesting their function as an umbrella for pollen grains during the rain events. We frequently observed intact flowers facing toward down-slope irrespective of slope direction like in forest-floor flowers on the slopes (Ushimaru et al. 2006) and often opened toward the sun direction in the study site, indicating no sun radiation preference and avoidance in the species (A. Ushimaru, personal observation). Flowering phenology of *P. grandiflorus* just overlaps rainy and typhoon seasons in Japan. Thus, horizontal orientation should have rainfall avoidance function in their flowers.

## Conclusion

In this study, our field experiment revealed that upward flowers cannot avoid damage from rainfall during the flowering period whereas downward flowers suffered from pollinator limitation in *P. grandiflorus*. Thus, horizontal flower orientation is suggested to evolve under both biotic and abiotic agents in this species. The very recent study suggested a similar adaptation in horizontally-orientated zygomorphic *Abelia* × *grandiflora* flowers (Yu et al., 2020). However, in the species, the role of horizontal orientation in rain protection is dubious because the anthers and stigmas are protuberant from the petals and always wet under rainy conditions (Yu et al., 2020). Thus, our results firstly demonstrated that horizontal orientation enhanced pollinator legitimate landings and female pollination success as well as protect mating-related organs, especially the anthers and secondary presented pollen grains in the male phase from rainfall damage in actinomorphic *P. grandiflorus* flowers. Because the adaptive significances of horizontal flower orientation in actinomorphic flowers with generalist pollination systems were examined only in this species, more other species should be investigated to generalize our findings.

## Supporting information

https://drive.google.com/file/d/1M06Qzx1BbB6RUfBmSl3RZWdW7fxj79jJ/view?usp=sharing

## ACKNOWLEDGEMENTS

We thank Gaku Hirayama, Fuma Kawakami, Akinari Sato, and Airi Asada for their help in field works and Koki R. Katsuhara for his suggestions on statistical analyses. We are grateful to the Sugadaira-kogen Hare snow resort for allowing our fieldworks. This research results were aided by a research fund (no. R1-3-17) to TN from Nagano Society for the promotion of science and a Grant-in-Aid for Scientific Research Programs (KAKENHI no. 19H03303) to AU from the Japan Society for the Promotion of Science.

## AUTHOR CONTRIBUTIONS

AU and RI firstly conceived the study and designed the methodology; TN, RI, YAY and AU collected the data; TN and AU analyzed the data and were involved in the writing of the manuscript. All authors contributed critically to the drafts and provided final approval for publication.

