## Supplementary material for "Horizontal orientation facilitates pollinator attraction and rain avoidance in radially symmetrical flowers": https://drive.google.com/file/d/1M06Qzx1BbB6RUfBmSl3RZWdW7fxj79jJ/view?usp=sharing

**TABLE S1.** Results of generalised linear mixed model (GLMM) and generalised linear model (GLM) analyses. Control flowers, male phase and distilled water (i.e., 0% sucrose solution) were used as the baselines for treatment, sex and sucrose concentration in the models (see text for details of the explanatory variables). Boldface indicates variables that had significant effects (*P* < 0.05).

| Response variable/ Explanatory variable | Estimated coefficient | S.E. | *z* value | *P* |
|---|---|---|---|---|
| Flower diameter / | | | | |
| **Intercept** | 4.699 | 0.110 | 42.480 | **<0.001** |
| **Sexual phase (female)** | 0.279 | 0.070 | 3.980 | **<0.001** |
| Pollinator Observation | | | | |
| All approaches per 20min. per flower / | | | | |
| Intercept | 0.387 | 0.212 | 1.820 | 0.068 |
| Treatment (Down) | -0.198 | 0.186 | -1.070 | 0.286 |
| Treatment (Up) | 0.178 | 0.173 | 1.030 | 0.303 |
| **Sexual phase (female)** | -0.426 | 0.205 | -2.080 | **0.037** |
| **Display size** | -0.291 | 0.094 | -3.080 | **0.002** |
| **Observation time** | 0.001 | 0.000 | 2.260 | **0.023** |
| Angle (Down)×Sex (female) | -0.074 | 0.333 | -0.220 | 0.822 |
| Angle (Up)×Sex (female) | -0.249 | 0.289 | -0.860 | 0.389 |
| All legitimate landings per 20min. per flower / | | | | |
| Intercept | -0.119 | 0.261 | -0.460 | 0.646 |
| **Treatment (Down)** | -0.819 | 0.240 | -3.410 | **<0.001** |
| Treatment (Up) | 0.135 | 0.200 | 0.680 | 0.496 |
| **Sexual phase (female)** | -0.833 | 0.267 | -3.120 | **0.001** |
| Display size | -0.211 | 0.118 | -1.790 | 0.074 |
| **Observation time** | 0.001 | 0.000 | 3.480 | **<0.001** |
| Angle (Down)×Sex (female) | -0.199 | 0.558 | -0.360 | 0.72 |
| Angle (Up)×Sex(female) | -0.274 | 0.383 | -0.720 | 0.473 |
| All petal landings per 20min. per flower / | | | | |
| **Intercept** | -1.483 | 0.715 | -2.070 | **0.038** |
| Treatment (Down) | -0.035 | 0.608 | -0.060 | 0.953 |
| Treatment (Up) | -0.526 | 0.652 | -0.810 | 0.42 |

| | | | | |
|---|---|---|---|---|
| Sexual phase (female) | -0.485 | 0.667 | -0.730 | 0.467 |
| Display size | -0.476 | 0.334 | -1.430 | 0.154 |
| Observation time | -0.002 | 0.001 | -1.880 | 0.059 |
| Treatment (Down)×Sex (female) | -0.054 | 1.001 | -0.050 | 0.956 |
| Treatment (Up)×Sex (female) | -0.074 | 1.092 | -0.070 | 0.946 |
| LM bee approaches per 20min. per flower / | | | | |
| Intercept | -0.327 | 0.251 | -1.300 | 0.193 |
| Treatment (Down) | -0.350 | 0.209 | -1.670 | 0.094 |
| Treatment (Up) | 0.046 | 0.194 | 0.240 | 0.813 |
| **Sexual phase (female)** | -0.739 | 0.289 | -2.550 | **0.011** |
| Display size | -0.179 | 0.103 | -1.740 | 0.082 |
| **Observation time** | 0.001 | 0.000 | 2.350 | **0.019** |
| Treatment (Down)×Sex (female) | 0.694 | 0.413 | 1.680 | 0.093 |
| Treatment (Up)×Sex (female) | 0.403 | 0.376 | 1.070 | 0.283 |
| LM bee legitimate landings per 20min. per flower / | | | | |
| Intercept | -0.580 | 0.301 | -1.950 | 0.051 |
| **Treatment (Down)** | -0.678 | 0.258 | -2.630 | **0.008** |
| Treatment (Up) | -0.015 | 0.223 | -0.070 | 0.944 |
| **Sexual phase (female)** | -1.318 | 0.394 | -3.340 | **<0.001** |
| Display size | -0.165 | 0.129 | -1.280 | 0.203 |
| **Observation time** | 0.001 | 0.001 | 2.660 | **0.007** |
| Treatment (Down)×Sex (female) | 0.386 | 0.669 | 0.580 | 0.563 |
| Treatment (Up)×Sex (female) | 0.605 | 0.499 | 1.210 | 0.224 |
| LM bee petal landings per 20min. per flower / | | | | |
| Intercept | -1.438 | 0.887 | -1.620 | 0.105 |
| Treatment (Down) | -1.139 | 0.816 | -1.400 | 0.163 |
| Treatment (Up) | -0.696 | 0.719 | -0.970 | 0.333 |
| Sexual phase (female) | -10.441 | 93.596 | -0.110 | 0.911 |
| Display size | -0.319 | 0.387 | -0.820 | 0.409 |
| Observation time | -0.005 | 0.002 | -1.870 | 0.062 |
| Treatment (Down)×Sex (female) | 11.192 | 93.601 | 0.120 | 0.905 |
| Treatment (Up)×Sex (female) | 1.081 | 120.080 | 0.010 | 0.993 |

| | | | | |
|---|---|---|---|---|
| S bee approaches per 20min. per flower / | | | | |
| Intercept | -0.860 | 0.450 | -1.910 | 0.056 |
| Treatment (Down) | -0.402 | 0.405 | -0.990 | 0.32 |
| Treatment (Up) | 0.407 | 0.350 | 1.160 | 0.245 |
| Sexual phase (female) | 0.014 | 0.383 | 0.040 | 0.97 |
| **Display size** | -0.678 | 0.222 | -3.050 | **0.002** |
| **Observation time** | 0.001 | 0.001 | 2.580 | **0.009** |
| Treatment (Down)×Sex (female) | -0.678 | 0.734 | -0.830 | 0.404 |
| **Treatment (Up)×Sex (female)** | -1.656 | 0.668 | -2.480 | **0.013** |
| S bee legitimate landings/ per 20min. per flower / | | | | |
| **Intercept** | -1.422 | 0.569 | -2.500 | **0.012** |
| **Treatment (Down)** | -1.679 | 0.620 | -2.710 | **0.006** |
| Treatment (Up) | 0.119 | 0.410 | 0.290 | 0.77 |
| Sexual phase (female) | -0.388 | 0.473 | -0.820 | 0.412 |
| **Display size** | -0.696 | 0.292 | -2.380 | **0.017** |
| **Observation time** | 0.003 | 0.001 | 3.650 | **<0.001** |
| Treatment (Down)×Sex (female) | -0.117 | 1.272 | -0.090 | 0.926 |
| Treatment (Up)×Sex (female) | -1.541 | 0.900 | -1.710 | 0.087 |
| S bee petal landings/ per 20min. per flower / | | | | |
| **Intercept** | -7.080 | 3.280 | -2.160 | **0.031** |
| Treatment (Down) | 0.421 | 1.800 | 0.230 | 0.816 |
| Treatment (Up) | 0.148 | 1.890 | 0.080 | 0.937 |
| Sexual phase (female) | 0.718 | 1.760 | 0.410 | 0.684 |
| Display size | -0.371 | 1.060 | -0.350 | 0.727 |
| Observation time | 0.002 | 0.002 | 0.880 | 0.381 |
| Treatment (Down)×Sex (female) | -1.140 | 2.360 | -0.480 | 0.629 |
| Treatment (Up)×Sex (female) | -17.000 | 0.002 | -0.010 | 0.995 |
| Syrphid fly approaches/ per 20min. per flower / | | | | |
| **Intercept** | -0.945 | 0.445 | -2.120 | **0.034** |
| Treatment (Down) | 0.375 | 0.334 | 1.120 | 0.261 |
| Treatment (Up) | 0.046 | 0.361 | 0.130 | 0.898 |

| | | | | |
|---|---|---|---|---|
| Sexual phase | -0.058 | 0.426 | -0.140 | 0.891 |
| Display size | -0.329 | 0.176 | -1.870 | 0.062 |
| Observation time | -0.001 | 0.009 | -1.810 | 0.07 |
| Treatment (Down)×Sex (female) | -1.572 | 0.852 | -1.840 | 0.065 |
| Treatment (Up)×Sex (female) | -0.444 | 0.646 | -0.690 | 0.492 |
| Syrphid fly legitimate landings/ per 20min. per flower / | | | | |
| **Intercept** | -2.800 | 0.855 | -3.280 | **0.001** |
| Treatment (Down) | -0.123 | 0.674 | -0.180 | 0.855 |
| Treatment (Up) | 0.718 | 0.598 | 1.200 | 0.23 |
| Sexual phase (female) | 0.403 | 0.657 | 0.610 | 0.54 |
| Display size | 0.061 | 0.317 | 0.200 | 0.845 |
| Observation time | -0.001 | 0.001 | -1.190 | 0.236 |
| Treatment (Down)×Sex (female) | -21.200 | 22400.000 | 0.000 | 0.999 |
| Treatment (Up)×Sex (female) | -2.460 | 1.260 | -1.950 | 0.051 |
| Syrphid fly petal landings/ per 20min. per flower / | | | | |
| **Intercept** | -2.870 | 1.160 | -2.460 | **0.014** |
| Treatment (Down) | 0.860 | 0.929 | 0.930 | 0.355 |
| Treatment (Up) | -0.815 | 1.290 | -0.630 | 0.527 |
| Sexual phase (female) | -0.400 | 1.290 | -0.310 | 0.757 |
| Display size | -3.220 | 0.485 | -0.660 | 0.506 |
| Observation time | 0.004 | 0.002 | -1.720 | 0.085 |
| Treatment (Down)×Sex (female) | -18.900 | 10400.000 | 0.000 | 0.999 |
| Treatment (Up)×Sex (female) | 1.510 | 1.820 | 0.830 | 0.407 |
| Pollen transfer success | | | | |
| Number of pollen grains (female) / | | | | |
| **Intercept** | 4.214 | 0.027 | 155.020 | **<0.001** |
| **Treatment (Down)** | -0.816 | 0.049 | -16.630 | **<0.001** |
| **Treatment (Up)** | -0.448 | 0.044 | -10.130 | **<0.001** |
| Number of pollen removal (male) / | | | | |
| **Intercept** | 12.100 | 0.001 | 22883.900 | **<0.001** |

| | | | | |
|---|---|---|---|---|
| **Treatment (Down)** | 0.023 | 0.001 | 31.330 | **<0.001** |
| **Treatment (Up)** | 0.011 | 0.001 | 15.430 | **<0.001** |
| Pollen germination experiments | | | | |
| Ratio of burst pollen grains (gaussian) / | | | | |
| **Intercept** | -1.053 | 0.248 | -4.250 | **<0.001** |
| **Sucrose concentration (5 %)** | -3.333 | 0.368 | -9.060 | **<0.001** |
| **Sucrose concentration (10%)** | -4.483 | 0.472 | -9.500 | **<0.001** |
| **Sucrose concentration (15%)** | -4.050 | 0.416 | -9.750 | **<0.001** |
| Sucrose concentration (20%) | -25.004 | 9080.300 | 0.000 | 1 |
| Ratio of germination (gaussian) / | | | | |
| **Intercept** | -3.909 | 0.277 | -14.100 | **<0.001** |
| **Sucrose concentration (5%)** | 3.365 | 0.297 | 11.300 | **<0.001** |
| **Sucrose concentration (10%)** | 3.317 | 0.296 | 11.200 | **<0.001** |
| **Sucrose concentration (15%)** | 3.367 | 0.297 | 11.300 | **<0.001** |
| **Sucrose concentration (20%)** | 3.426 | 0.298 | 11.500 | **<0.001** |

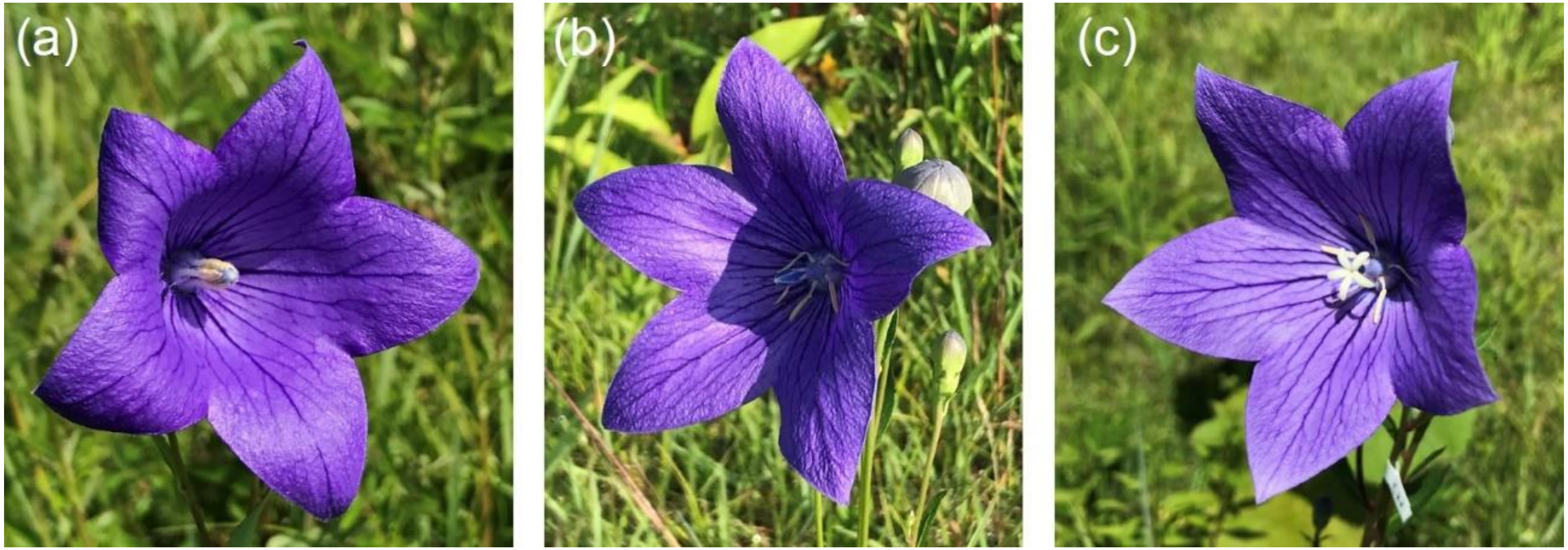

**FIGURES S1.** Photographs of a flower soon after opening (a) and those in the male- (b) and female-phases (c) of *Platycodon grandifloras*.

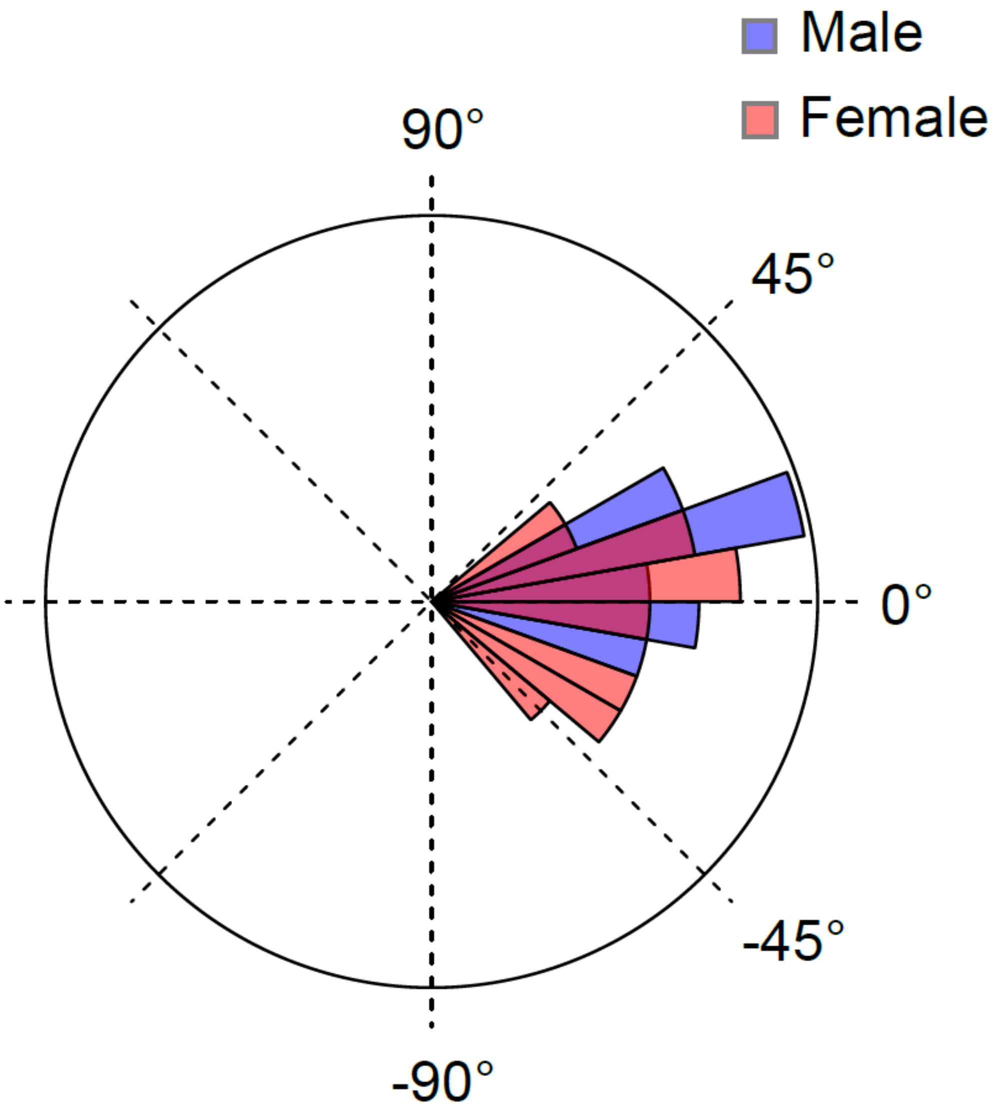

**FIGURE S2.** Circular histograms of flower angles of both male- (blue bars) and female-phase (red bars) flowers in 2021. Flower angle did not significantly differ between male- and female-phase flowers ($\chi^2$ =7.771, $P$ = 0.169)

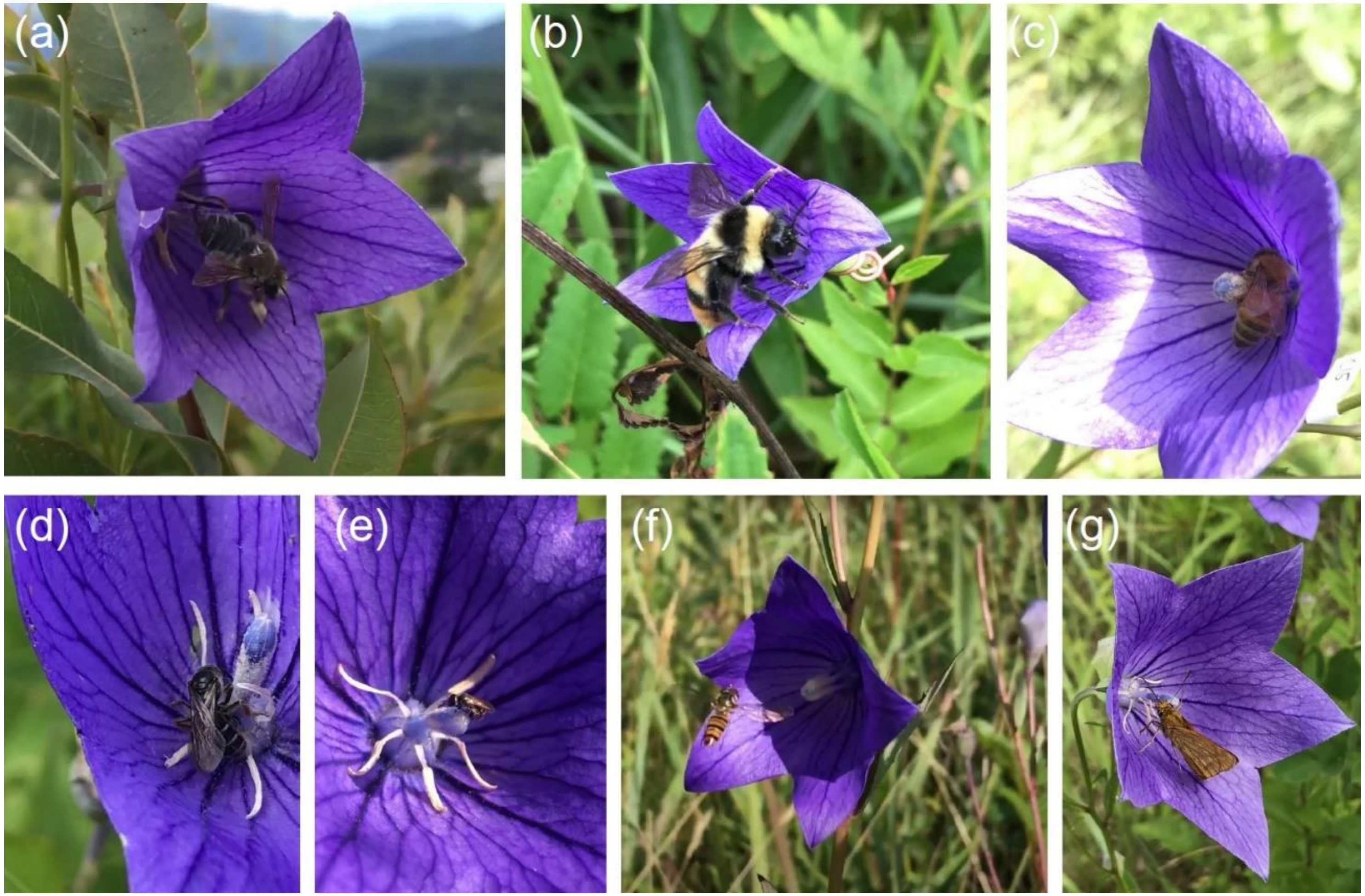

**FIGURE S3.** Diverse pollinators on *Platycodon grandiflorus*: large and medium bees, *Megachile* sp. (a), *Bombus hypocrita* (b), *Apis mellifera* (c); small bees; *Andrena* sp. (d), Halictidae sp. (e); a syrphid fly, *Episyrphus balteatus* (f); a skipper, *Ochlodes ochraceus* (g).

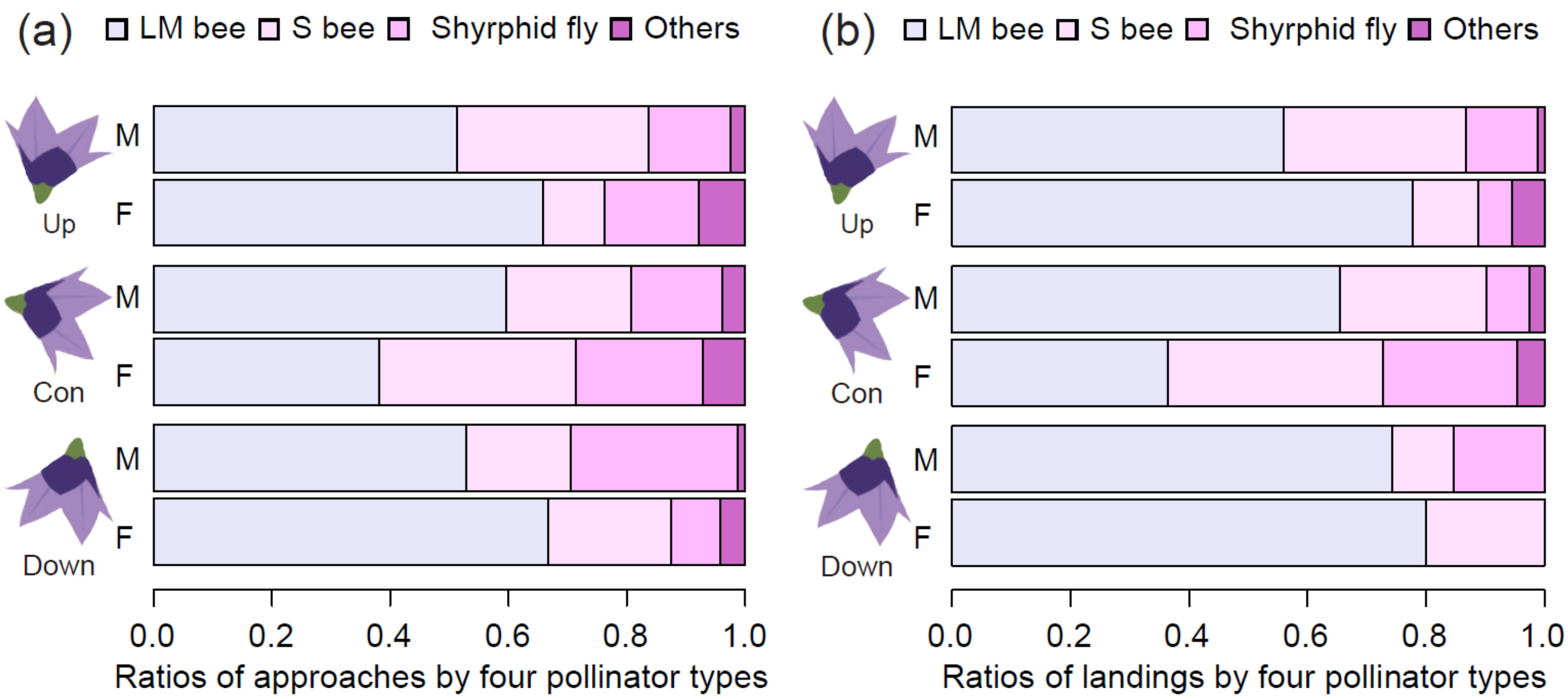

**FIGURE S4.** The ratios of approaches (a) and legitimate landings (b) by large and medium bees (LM bee, lavender), small bees (S bee, thistle), syrphid flies (plum) and other pollinators (orchid) for the experimental (Con, Up and Down) flowers. The ratios of three dominant pollinator groups differed significantly among flower types and between sexual phases for approaches ($\chi^2$ = 22.534, $P$ = 0.012), but did not differ for legitimate landings ($\chi^2$ = 17.63, $P$ = 0.06).

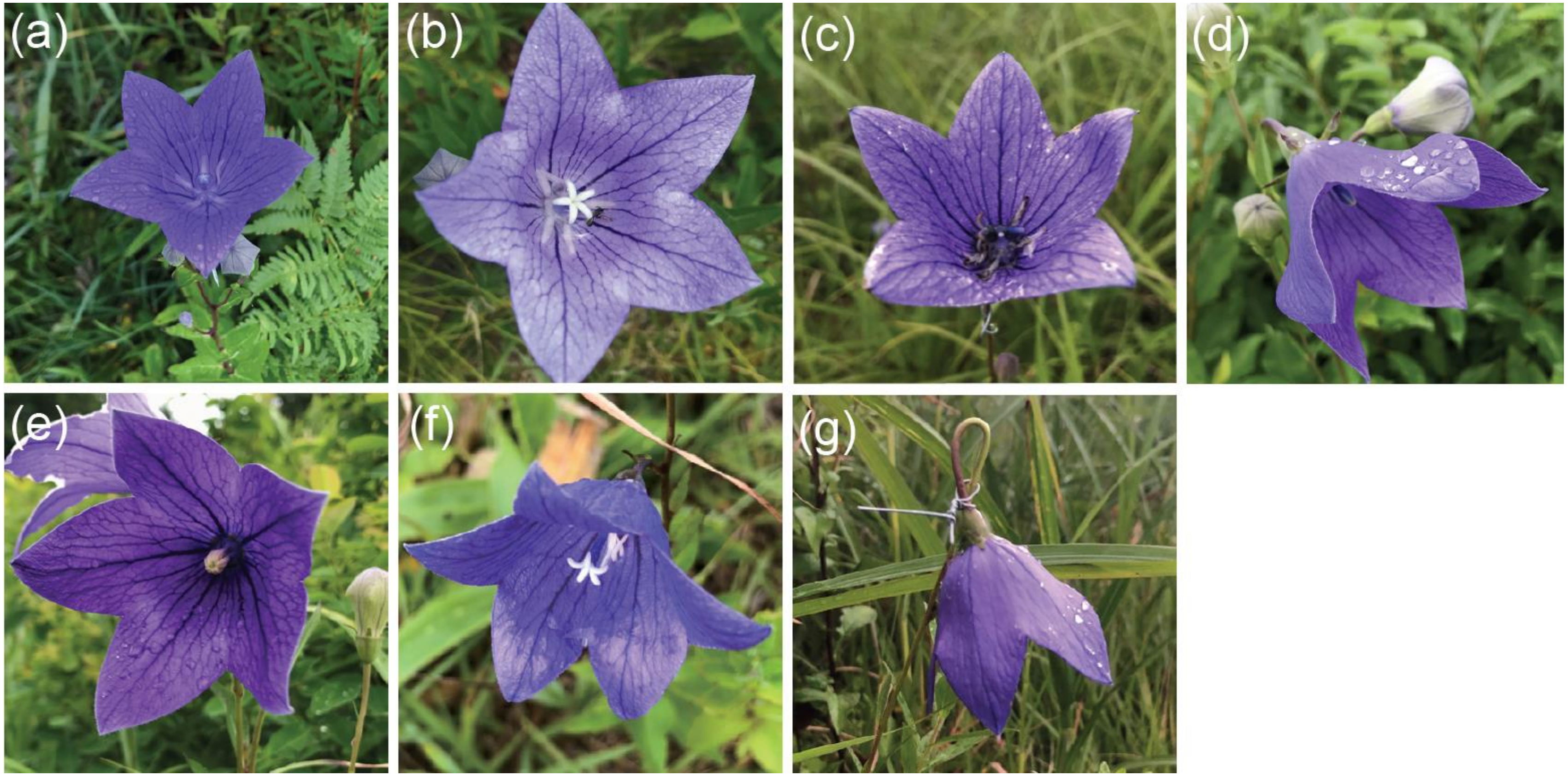

**FIGURE S5.** Photos of experimental and intact flowers of *Platycodon grandifloras* after rain (treatment, precipitation recorded in the observation day or the day before): pistil hairs and anthers of male phase flower was soaked (a: Up, 3.5 mm/d), stigma of female phase flower and a pollinator were soaked (b: Up, 3.5 mm/d), the stigma of female phase flower was broken (c: Up, 6.0 mm/d), horizontal and dowbward flowers protected the anthers, stigma and pollen from rainfall (d: Con; intact, 10.0 mm/d; e: intact, 6.0 mm /d, f:intact, 2 mm/d; g: Down, 6.0 mm/d).

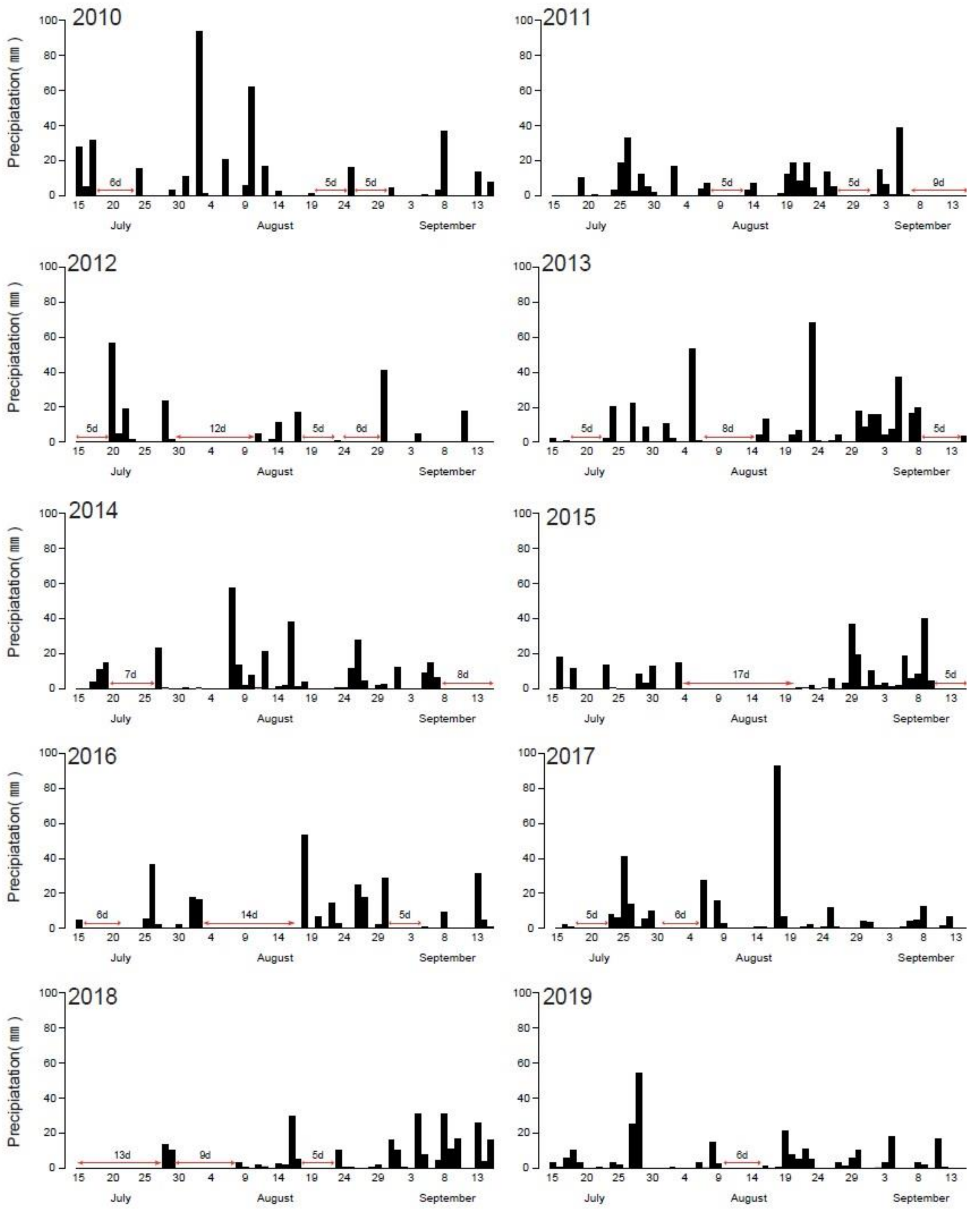

**FIGURE S6.** Precipitation of the flowering season which were observed by Automated Meteorological Data Acquisition System in Sugadaira, Nagano prefecture (N36°31’9, E138°19’5).
